# Long Read Annotation (LoReAn): automated eukaryotic genome annotation based on long-read cDNA sequencing

**DOI:** 10.1101/230359

**Authors:** David E. Cook, Jose Espejo Valle-Inclan, Alije Pajoro, Hanna Rovenich, Bart PHJ Thomma, Luigi Faino

## Abstract

Single-molecule full-length cDNA sequencing can aid genome annotation by revealing transcript structure and alternative splice-forms, yet current annotation pipelines do not incorporate such information. Here we present LoReAn (<u>Lo</u>ng <u>Re</u>ad <u>An</u>notation) software, an automated annotation pipeline utilizing short- and long-read cDNA sequencing, protein evidence, and *ab initio* prediction to generate accurate genome annotations. Based on annotations of two fungal and two plant genomes, we show that LoReAn outperforms popular annotation pipelines by integrating single-molecule cDNA sequencing data generated from either the PacBio or MinION sequencing platforms, and correctly predicting gene structure and capturing genes missed by other annotation pipelines.

## Background

Genome sequencing has advance nearly every discipline within the biological sciences, as the ongoing decreasing sequencing costs and increasing computational capacity allows many laboratories to pursue genomics-based answers to biological questions. New sequencing technologies designed to sequence longer contiguous DNA molecules, such as Pacific Biosciences’ (PacBio) Single Molecule Real Time sequencing (SMRT) and Oxford Nanopore Technologies’ (ONT) MinION, have ushered the most recent genomics revolution [1]. These advances are further enhancing the ability to generate high-quality genome assemblies of large, complex eukaryotic genomes [2-5].

A high-quality genome assembly, represented by (near-)chromosome completion, can help to address many biological questions, but often requires functional features to be further defined [6]. The process of genome annotation, i.e. the identification of protein-coding genes and their structural features such as intron-exons boundaries, is important to capture biological values of a genome assembly [7]. Genomes can be annotated using computer algorithms in so-called *ab initio* gene predictions, as well as using wet-lab generated data, such as cDNA or protein datasets for evidence-based predictions, and current annotation pipelines typically incorporate both types of data g[7, 8]. *Ab initio* gene prediction tools are based on statistical models, most often Hidden Markov Models (HMMs), that are trained using known proteins, and typically perform well at predicting conserved or core genes [7, 9]. However, the *ab initio* prediction accuracy decreases for organism-specific genes, for genes encoding small proteins and those containing introns in untranslated regions (UTRs). Furthermore, *ab initio* annotation of non-model genomes remains challenging as appropriate training data is not always available. To improve genome annotations, cDNA sequencing (RNA-seq) data can be incorporated to train *ab initio* software [10] and to provide additional evidence for defining accurate gene models [11]. However, it remains challenging to annotate a genome with short-read RNA-seq data due to difficulties in unequivocally mapping these reads, and because single reads do not span a gene’s full length. Consequently, the coding structure must be computational inferred.

Current annotation pipelines use a combination of *ab initio* and evidence based predictions to generate accurate consensus annotations. MAKER2 is a user-friendly, fully automated annotation pipeline that incorporates multiple sources of gene prediction information and has been extensively used to annotate eukaryotic genomes [12-16]. The <u>B</u>road Institute Eukaryotic Genome <u>A</u>nnotation <u>P</u>ipeline (here referred to as BAP) has mainly been used to annotate fungal genomes [17-19] and integrates multiple programs and evidences for genome annotation [20, 21]. CodingQuarry is another gene prediction software that utilizes general HMMs for gene prediction using both RNA-seq data and genome sequence [22]. A limitation of these annotation pipelines is that they give little weight to experimental evidence such as short read RNA-seq and cannot exploit gene structure information from single-molecule cDNA sequencing.

In addition to improving the genome assembly [23], long-read sequencing data can be used to improve genome annotation. The use of single-molecule cDNA sequencing can increase the accuracy of automated genome annotation by improving genome mapping of sequencing data, correctly identifying intron exon boundaries, directly identifying alternatively spliced transcripts, identifying transcription start and end sites, and providing precise strand orientation to single exons genes [24-26]. However, several hurdles limit the implementation of long-read sequencing data into automated genome annotation, such as the higher per-base costs when compared to short-read data, the relatively high error rates for long-read sequencing technologies, and the lack of bioinformatics tools to integrate long-read data into current annotation pipelines [27, 28]. The first two limitations are addressed by the continual reduction in sequencing cost and improving base calling by long-read sequencing providers, and the development of bioinformatics methods to correct for sequencing errors [29, 30]. To address the disconnection between genome annotation pipelines and the latest sequencing technologies, we developed the <u>Lo</u>ng <u>Re</u>ad <u>An</u>notation (LoReAn) pipeline. LoReAn is an automated annotation pipeline that takes full advantage of MinION or PacBio SMRT long-read sequencing data in combination with protein evidence and *ab initio* gene predictions for full genome annotation. Short-read RNA-seq can be used in LoReAn to train *ab initio* software. Based on the reannotation of two fungal and two plant species, we demonstrate that LoReAn can provide annotations with increased accuracy by incorporating single-molecule cDNA sequencing data from different sequencing platforms.

## Results

### Long-read annotation (LoReAn) design and implementation

The LoReAn pipeline can be conceptualized in two phases. The first phase involves genome annotation based on *ab initio* and evidence-based predictions (Fig. 1a: blue arrows) and largely follows the workflow previously described in the BAP [20, 21]. This first phase produces a full-genome annotation and requires the minimum input of a reference genome, protein sequence of known and, possibly, related species, and a species name from the Augustus prediction software database [31]. Two changes were implemented into the first phase of LoReAn, which we refer to as BAP+. One alteration is that LoReAn uses RNA-seq reads as input in combination with the BRAKER1 software [10] to produce a species specific database for the Augustus prediction software. Additionally, RNA-seq data is assembled into full-length cDNA using Trinity software [32] and the assembled transcripts are aligned to the genome using both PASA [20] and GMAP [33]. The output of PASA software is passed to Evidence Modeler (EVM) [20] as cDNA evidence while the output of GMAP is given to EVM as *ab initio* software. GMAP output passed as *ab initio*-evidence guarantees that genes not predicted by *ab initio* software like Augustus and GeneMark but present in the transcriptome are passed to Evidence Modeler.

**Fig. 1.**
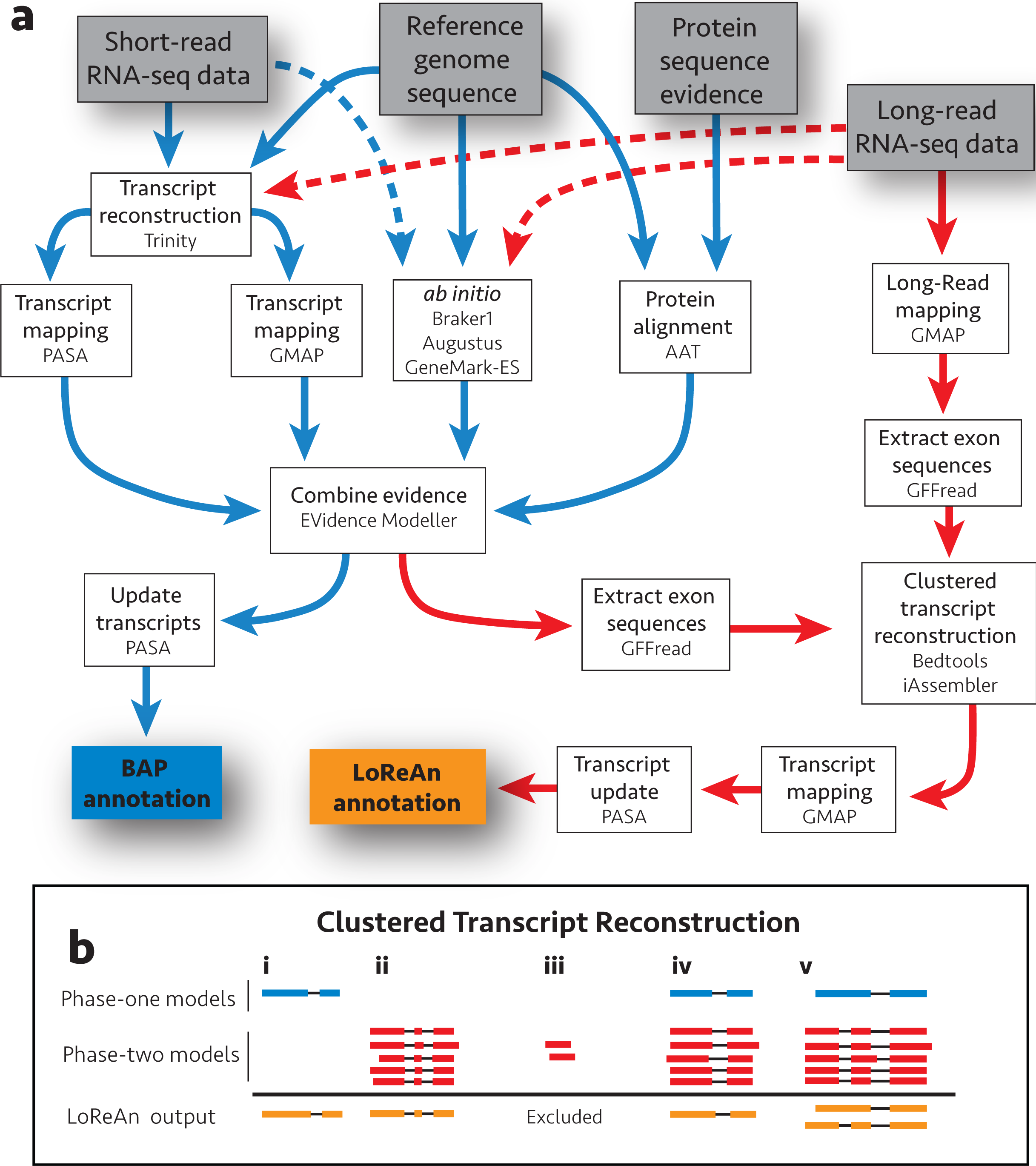
Schematic overview of the LoReAn pipeline and clustered transcript reconstruction. **a** Illustration of the computational workflow for the LoReAn pipeline. Grey boxes represent input data and each white box represents a step in the annotation process with mention of the specific software. The boxes connected by blue arrows integrate the steps from the previously described BAP [20]. The LoReAn pipeline (boxes connected by red arrows) integrates the BAP workflow, but additionally incorporates long-read sequencing data. The orange box, ‘Final BAP annotation’ represents the annotation results from the BAP pipeline used for comparison in this study. Dashed arrows represent optional steps for the pipeline. **b** Illustration of the clustered transcript reconstruction. Gene models are depicted as exons (boxes) and connecting introns (lines). Blue models represent BAP annotations, while red models represent hypothetical long-reads mapped to the genome. Orange models represent consensus annotations reported in the final LoReAn output. Various scenarios can occur: i: High confidence predictions from the BAP are kept regardless of whether they are supported by long-reads. ii & iii: Clusters of mapped long-reads are used to generate a consensus prediction model, unless the model is supported by less than a user-defined minimum depth. iv: Overlapping BAP and mapped long-reads are combined to a consensus model. v: Two annotations are reported if no consensus can be reached for the BAP and clustered long-read data.

The second phase of LoReAn incorporates single-molecule cDNA sequencing with the annotation results of the first phase by utilizing a novel approach to reconstruct full-length transcripts (Fig. 1a: red arrows). Singlemolecule long-read sequencing reads are mapped to the genome using GMAP, which allows the determination of transcript structure (i.e. start, stop and exon boundaries) from a single cDNA molecule [34]. The underlying reference sequence is extracted to overcome sequence errors associated with long-read sequencing, and these sequences are combined with the gene models from the first phase in a process we refer to as ‘clustered transcript reconstruction’ (Fig 1a and b). Through this process, consensus gene models are built by combining the first and second phase gene models that cluster at the same locus. Optionally, model clustering can be done in a strand-specific manner (LoReAn stranded, main text in Additional file 1 for details) where only gene models mapping on the same DNA coding strand are used to build a consensus model. These high-confidence models are mapped back to the reference using GMAP to correct open reading frames and subsequently, PASA is used to update the gene models by identifying untranslated regions (UTRs) and alternatively spliced transcripts to generate a final annotation. Sequence-based support for the final gene models (Fig. 1b orange models) can come from the first phase annotation alone (Fig. 1b i), the second phase given a sufficient level of support (Fig. 1b ii, iii), or through a combination of the two phases (Fig. 1b iv, v). If a single consensus annotation cannot be reached between the two phases, both annotations are kept in the final output (Fig. 1b v).

### LoReAn produces the highest accuracy gene predictions

To test the performance of LoReAn, we re-annotated the genome sequence of the haploid fungus *Verticillium dahliae,* an important pathogen of hundreds of plant species including many crops [35, 36]. The genome of *V. dahliae* strain JR2 was used for testing LoReAn because it is assembled into complete chromosomes and has a manually curated annotation, providing a high-confidence resource for reference [2]. The output of 54 annotations were compared, of which 24 were produced using LoReAn, 12 using BAP and 12 using BAP+ with different genome masking and *ab initio* options (description in Additional file 1), along with output from the annotation software MAKER2, CodingQuarry, BRAKER1, Augustus and two from GeneMark-ES (Fig 2a; Additional file 2: Table S1). The quality of the annotation outputs were determined by comparing each to the reference annotation for exact matches to either genes, transcripts or exon locations. These comparisons were used to calculate sensitivity (how much of the reference is correctly predicted), specificity (how much of the prediction is in the reference), and accuracy (an average of sensitivity and specificity). We calculated these metrics based on commonly described methods used within the gene prediction community (see methods and references [7, 37, 38]). Genome masking prior to annotation significantly affected the accuracy of predicted gene models, with partially masked or non-masked genome inputs producing the most accurate annotations (Fig 2a; Additional file 1: Fig S1a - S3a, Additional file 2: Table S2). On average, the ‘fungus’ option of the *ab initio* software GeneMark-ES produced the most accurate gene, transcript, and exon predictions (Fig 2a; Additional file 1: Fig S1b - S3b; Additional file 2: Table S2 - S4). Gene predictions from LoReAn using coding strand information (LoReAn-s) had the highest accuracy across the tested conditions for exact match genes to the reference annotation (Fig. 2a; Table 1).

**Fig. 2.**
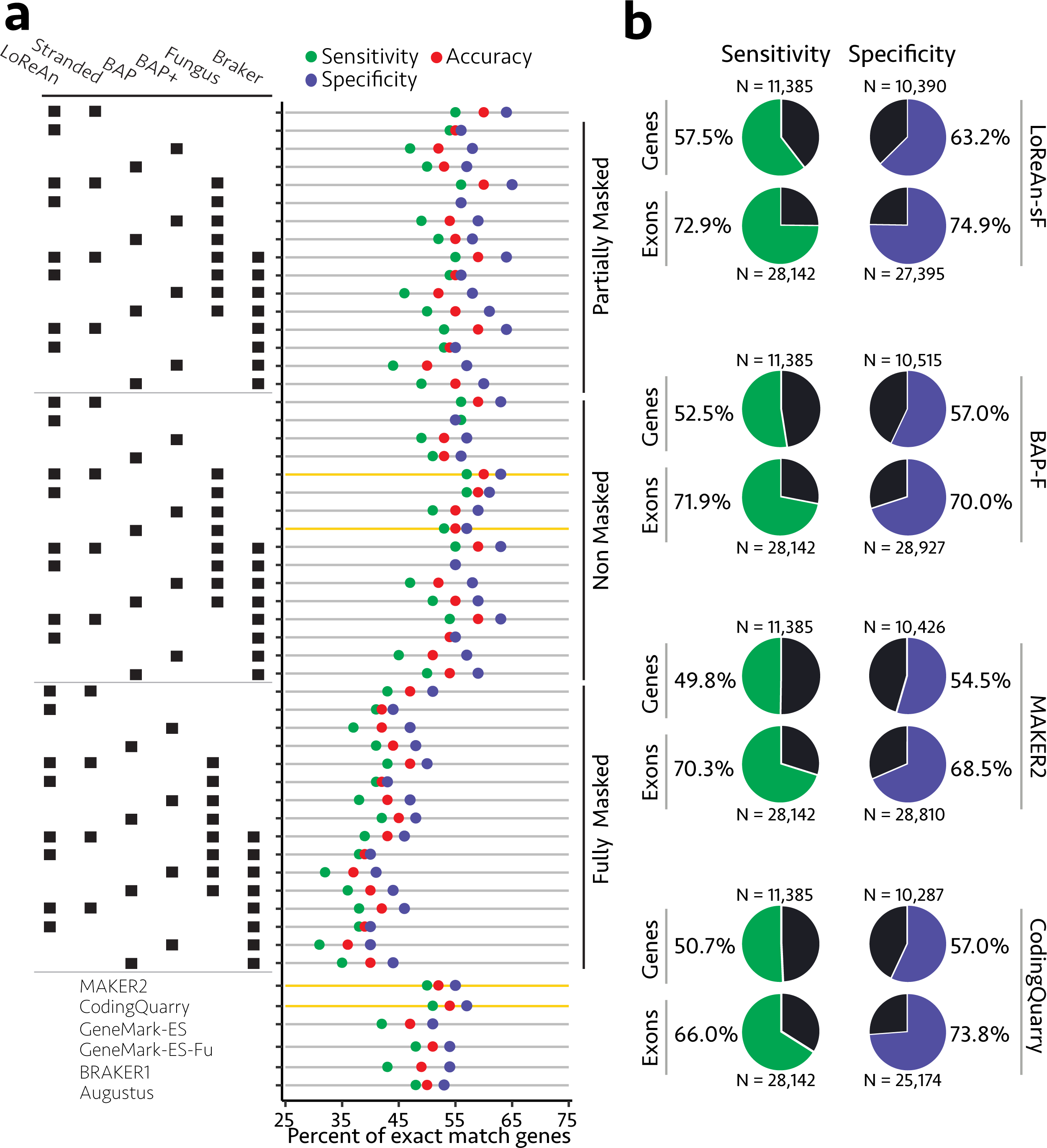
Annotation quality summary for exact match genes to the reference. **a** Each horizontal bar represents an annotation output, and each colored dot represents the sensitivity (green), specificity (purple) and accuracy (red). The annotations are labelled using the left grid table, where the group of horizontal black dots defines the parameters used in the annotation. Possible parameters include using the LoReAn, BAP or BAP+ pipeline, stranded mode for LoReAn (Stranded), the fungus option for GeneMark-ES (Fungus), or the BRAKER1 program for Augustus (BRAKER1). Each set of 16 annotations are grouped by the level of reference masking, Partially Masked, Non-Masked or Fully Masked (right label). The results from additionally tested annotation pipelines are shown at the bottom. The four annotations highlighted with a yellow horizontal bar were used for subsequent analysis. **b** Sensitivity and specificity for exact match genes and exons for the best annotations highlighted in yellow in a. For the sensitivity column, the number N represents the number of reference features, and the green sector of the pie chart shows the sensitivity. For example, the top sensitivity chart indicates that the LoReAn-sF pipeline annotated 57.5% (6,546) of the reference annotations 11,385 genes with exact feature matches.

A single output from LoReAn, BAP, MAKER2 and CodingQuarry were selected for in-depth comparison (Fig. 2a, horizontal lines highlighted in yellow; Fig. 2b). The LoReAn-stranded run using the ‘fungus’ option of GeneMark-ES (referred to as LoReAn-sF throughout) and the BAP run using the fungus option of GeneMark-ES (referred to as BAP-F throughout) using a non-masked genome as input were selected because they had the highest accuracy and used similar settings, thereby enabling comparisons (Additional file 2: Table S1). Default settings for MAKER2 and CodingQuarry were run with as similar input to the LoReAn and BAP pipelines as possible. The LoReAn-sF output had the highest gene and exon sensitivity and specificity compared to the other three pipelines, showing a 13% increase in gene sensitivity and 9% increase in gene specificity compared to the next best performing pipeline, BAP-F (Fig. 2b).

Collectively, the results from testing gene prediction options and pipelines show that genome masking prior to annotation and *ab initio* options can impact the quality of a genome annotation. Across the tested settings, the LoReAn pipeline produces the highest quality gene predictions when compared to the reference annotation. Overall, LoReAn-stranded produced the best annotation predictions, highlighting that incorporating single-molecule cDNA information in the annotation process significantly improves the output.

**Table 1:** Annotation quality metrics for exact match genes for the tested pipelines

| Pipeline<sup>c</sup> | Sensitivity<sup>b</sup> | Specificity<sup>b</sup> | Accuracy<sup>b</sup> |
| --- | --- | --- | --- |
| BAP<sup>a</sup> | 50.0% | 60.5% | 55.3% |
| BAP+<sup>a</sup> | 50.7% | 58.7% | 54.7% |
| LoReAn<sup>a</sup> | 57.3% | 61.0% | 59.2% |
| LoReAn-s<sup>a</sup> | 57.5% | 63.2% | 60.3% |
| MAKER2 | 49.8% | 54.5% | 52.1% |
| CodingQuarry | 50.7% | 57.0% | 53.8% |
| Augustus | 47.8% | 53.0% | 50.4% |
| BRAKER1 | 43.4% | 54.4% | 49.9% |
| GeneMark-ES | 42.4% | 50.7% | 46.6% |
| GeneMark-ES+Fungus | 47.5% | 54.4% | 51.0% |

<sup>a</sup>Results from the highest quality annotation are shown
<sup>b</sup>Each quality metric was calculated against the *V. dahliae* strain JR2 reference. Details on how each was calculated and their definitions can be found in the Methods section.
<sup>c</sup>Pipelines: BAP - Broad Annotation Pipeline; BAP+ - modified BAP; LoR\_M - LoReAn using masked input genome; LoR - LoReAn; LoR\_S\_M - LoReAn stranded mode using masked input genome; LoR\_S - LoReAn stranded mode.

### LoReAn predicts the greatest number of high-confidence genes compared to other pipelines

The four best gene predictions; LoReAn-sF, BAP-F, MAKER2 and CodingQuarry, were compared head-to-head in the absence of a reference annotation to determine differences in gene prediction. There were 4,584 genes with the same predicted structure (i.e. start, stop, intron position) from the 4 pipelines, equivalent to approximately 40% of the genes in the reference annotation (Fig. 3a). BAP predicted the fewest unique genes (1,352), while MAKER2 predicted the most (3,157) (Fig. 3a). However, the use of exact match gene structure to identify unique coding sequence is potentially misleading, as two gene predictions can code for the same or a similar protein without the exact same structure. To generate a more biologically relevant comparison of unique protein coding differences, we grouped translated protein sequences of each annotation into homologous groups using orthoMCL [39, 40]. Using these groups, we identified protein coding sequences that were unique to a single annotation pipeline, referred to as singletons. We identified 1,429 singletons across the four annotations, with CodingQuarry predicting the most (461) and BAP-F predicting the fewest (180). The validity of the singletons were analyzed by checking their support from short-read RNA-seq data. Coding sequences from the LoReAn-sF protein singletons averaged 80% coverage across the predicted gene model’s length, statistically significantly greater than the singleton coverage from the other pipelines (Fig. 3b). The log2 length of the singletons did not significantly change across the LoReAn-sF, BAP-F and MAKER2 results (Fig. 3c). Additionally, we checked the singletons for introns, and grouped them by RNA-seq coverage, as genes with introns and RNA-seq support are more likely to be true genes. Singletons that contain at least one intron and have RNA-seq reads covering at least 75% of their length were considered the highest-confidence models. The LoReAn-sF pipeline had the greatest number of singletons in this high-confidence category, 241, which represents 55.1% of the total singletons predicted by the pipeline. MAKER2 also predicted many singletons in this category, 176, which was 50.1% of the singletons predicted by the pipeline (Fig. 3d, green wedge). In contrast, the CodingQuarry and BAP-F pipelines predicted the most low-confidence singletons, those with no introns and lower RNA-seq support, representing a greater proportion of the singletons predicted by the pipelines (Fig. 3d). For research projects aimed at identifying new protein coding genes, these results suggest the LoReAn-sF pipeline offers the greatest chance at identifying novel, high-confidence protein coding genes.

**Fig. 3.**
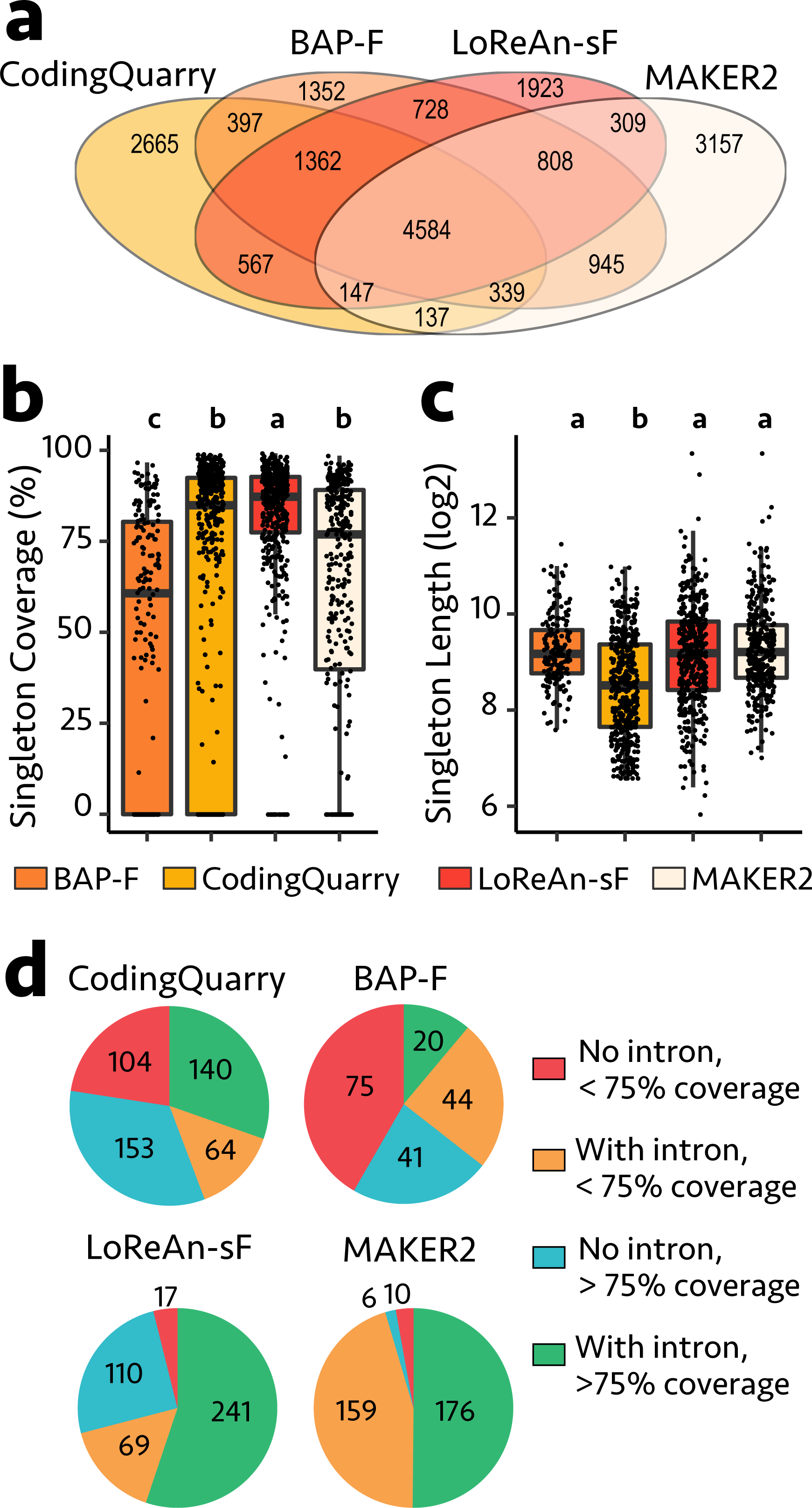
Comparison of the unique genes annotated from each of the four pipelines. **a** To directly compare the annotation output from the four pipelines against each other, we identified the number of exact match genes across the four annotations. The Venn diagram shows that 4,646 genes were annotated with the exact same features across all four pipelines. The numbers captured by only a single annotation pipeline are considered singletons-genes whose structure is uniquely annotated by a given pipeline. Note, these singletons do not necessarily represent unique loci. **b** The percent length of each gene model covered by RNA-sequencing data is shown as a bar chart for each annotation pipeline. Each box plot represents the standard interquartile ranges and each dot represents a data point. An ANOVA was calculated for each metric, such as singleton coverage ~ pipeline, and post-hoc tested using Tukey Honestly Significant Difference (HSD) with alpha = 0.05. Letters shown above each box plot represents the HSD groupings. **c** Same as in b except the lengths of each predicted model were analyzed as log2 values. **d** The orthoMCL singletons from each pipeline were grouped into one of four categories shown in the key representing if the singleton contained an intron or not and if the singleton’s length was covered by over 75% with RNA-seq data. The number of singletons within each of the four categories is shown.

### LoReAn gene predictions are the most accurate based on reference independent analysis

To evaluate the annotation output in the absence of a reference, we devised an approach to quantify annotation accuracy based on empirical data. The locations of predicted introns from the annotation outputs were compared to the locations of the inferred introns from long- and short-read mapped data. This analysis shows that LoReAn outputs using non- or partially masked genomes have the highest exact match intron accuracy (Fig. 4, points closest to top right corner). To validate this approach, the exact match intron accuracy from mapped reads were correlated with the to exact match gene accuracy from the reference annotation. This analysis shows a significant positive correlation between the reference dependent and independent assessments (r = 0.88, p-value < 2.2e-16, spearman correlation) (Additional file 1: Fig. S4). This indicates that the empirical annotation assessment is an alternative method to assess gene prediction accuracy in the absence of an annotation or the absence of a high-confidence annotation.

**Fig. 4.**
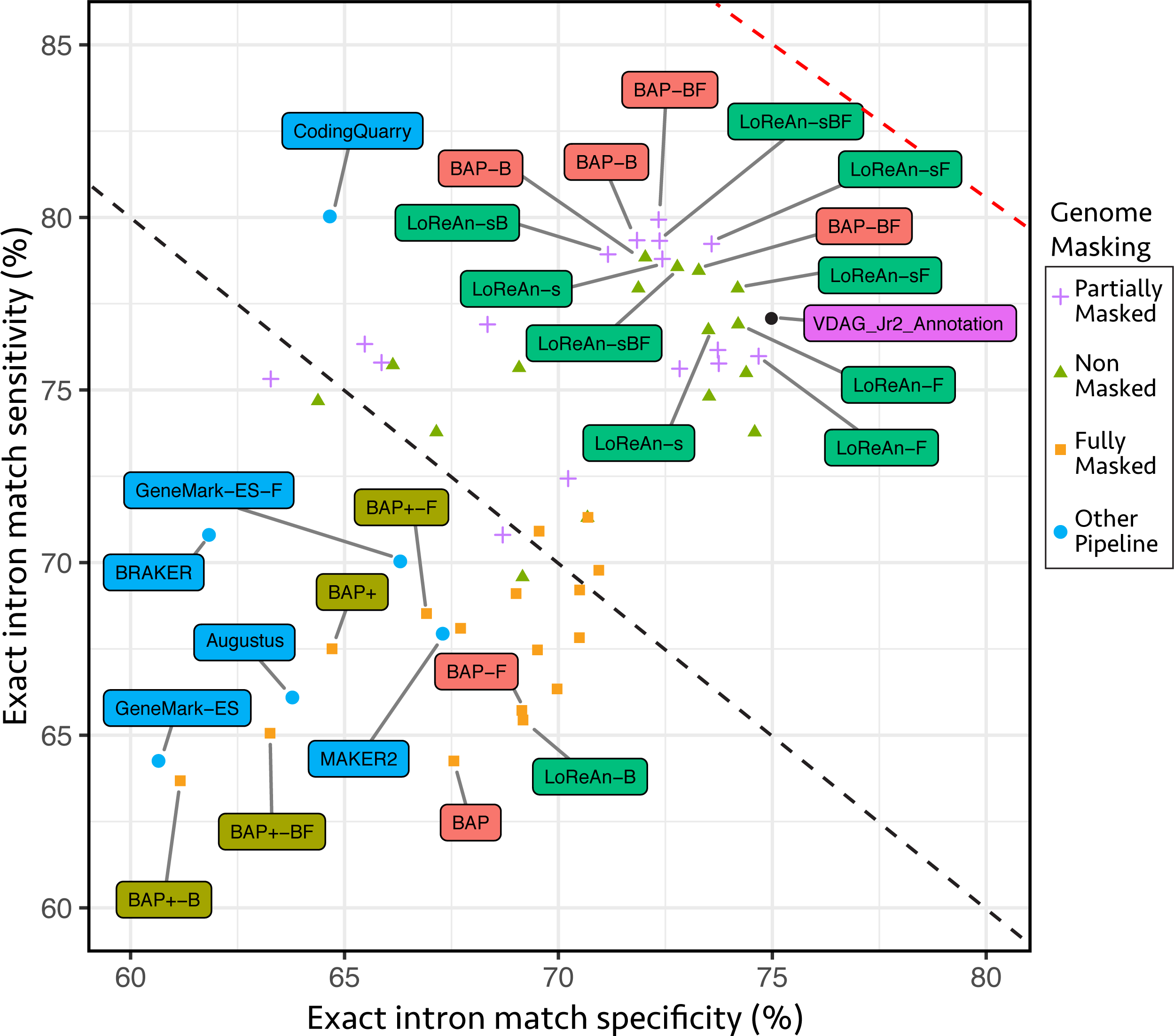
LoReAn gene predictions are the most accurate based on analysis of intron location. The quality of 55 gene predictions using the *V. dahliae* genome were assessed using exact intron matches Sensitivity (y-axis) and specificity (x-axis) were mapped, and the symbols represent their accuracy (average of sensitivity and specificity). The dashed black and red lines represent 70% and 80% accuracy respectively. Individual predictions with an accuracy greater than 75% or lower than 68%, along with the independent pipelines are labeled in colored boxes connected to their corresponding points with a grey line. The results of the *V. dahliae* JR2 strain annotation compared to the mapped introns is shown in black, labeled VDAG_Jr2_Annotation.v5. s — stranded; B — Braker1; F — Fungus option.

### Only the LoReAn pipeline correctly annotates the *Ave1* effector locus

Plant-pathogenic fungi encode *in planta-secreted* proteins, termed effectors, which serve to facilitate infection [41, 42]. Effectors are generally characterized as lineage-specific small, secreted, cysteine-rich proteins with generally no characterized protein domains or homology, characteristics which can make effectors difficult to predict with automated annotation [43]. To test how LoReAn and the other annotation pipelines performed at a specific effector locus, we detailed the annotation results for the *V. dahliae Ave1* locus, which encodes a small-secreted protein that functions to increase virulence during plant infection [44]. As previously reported, a considerable number of short RNA-seq reads uniquely map to the *Ave1* locus [44], along with single-molecule cDNA reads identified here (Fig. 5a). Interestingly, the MAKER2, BAP, and CodingQuarry pipelines, along with the Augustus and GeneMark-ES software fail to predict the previously characterized *Ave1* gene, despite the abundance of uniquely-mapped reads (Fig. 5b; Additional file 1: Fig. S5). Intriguingly, the MAKER2 and BAP pipelines predict a separate gene on the opposite strand located to the 3’ end of the *Ave1* gene that is absent in the reference annotation. The LoReAn-sF and BAP+ pipelines predict two genes at the locus, one corresponding to the known *Ave1* gene, and an additional gene to the 3’ end of *Ave1* (called *Ave1c*), similar to the gene model identified by MAKER2 and BAP (Fig. 5b; Additional file 1: Fig. S5).

**Fig. 5.**
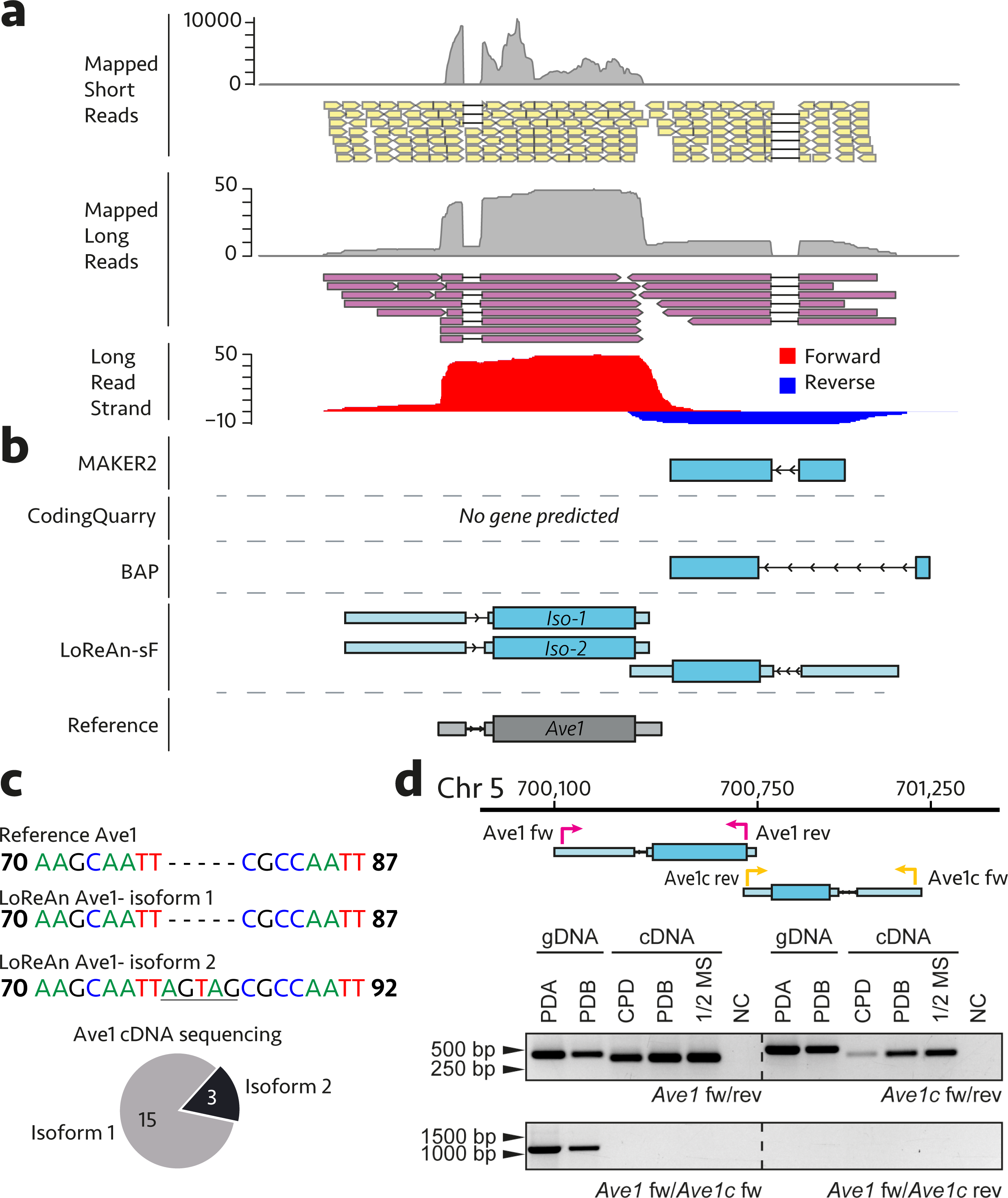
The LoReAn pipeline most accurately annotates a specific fungal locus encoding a strain specific gene. **a** Short-read RNA-seq data mapped to the locus are shown as a coverage plot (grey peaks) and as representative individual reads (yellow boxes). Long-reads from single-molecule cDNA data mapped to the locus are shown as a coverage plot (grey peaks) and representative reads (purple boxes). Think black lines linked mapped reads represent gaps in the mapped reads and are indicative of introns. The long-read data was split by mapping strand and coverage plots for forward (red) and reverse (blue) coverage plots. **b** Gene model predictions from the four annotation pipelines are illustrated. Light blue boxes represent untranslated regions (5’ and 3’ UTR), dark blue boxes represent coding sequence boundaries, and thin black lines depict introns. Arrows in the introns indicate the direction of transcription. The MAKER2 and BAP pipelines predict a single transcript coded on the reverse strand at the 3’ end of the known *Ave1* transcript. Coding Quarry does not predict a gene at the locus. LoReAn predicts two transcripts corresponding to the *Ave1* gene along with the similar transcript predicted by MAKER2 and BAP. The reference *Ave1* transcript is shown in grey. **c** To confirm the presence of an alternative splice site in the 5’UTR of the *Ave1* transcript, 18 cDNA clones were randomly chosen and sequenced. Isoform 1 sequence is identical to the reference *Ave1* sequence and was identified in 15 of the 18 clones. Isoform 2 has a 5 bp insertion in the 5’UTR resulting from an alternative exon splice site and was identified in 3 of the 18 sequenced clones. The *Ave1* reference sequence is shown from bases 71 through 86. **d** The presence of *Ave1* and the additional gene transcribed to the 3’ end of *Ave1,* termed *Ave1close(Ave1c),* was confirmed using PCR on gDNA and cDNA. PCR using gene specific primers, termed *Ave1* fw + rev (pink arrows) or *Ave1c* for + rev (yellow arrows), shows that both genes are expressed in either potato dextrose broth (PDB) Czapek-dox (CPD) or half-strength Murashige-Skoog (1/2MS) media. The inverse orientation of the two genes was confirmed using forward primers only, which amplified the entire locus resulting in a band of approximately 1,118 bp, but does not amplify product using cDNA as the template.

LoReAn-sF additionally predicts two mRNAs corresponding to the previously characterized *Ave1* gene, termed isoform-1 and -2 (Fig. 5b). To confirm the presence of two *Ave1* isoforms, cDNAs were amplified and cloned into vectors, and 18 clones were randomly selected for sequencing. A majority of the sequenced transcripts, 15 of 18, have a sequence corresponding to isoform-1, the known *Ave1* transcript, while the other 3 were the isoform-2 sequence (Fig. 5c). The isoform-2 transcript is the result of an alternative splice junction 5 bp upstream of the previously identified splice site in the *Ave1* 5’ UTR intron, and is not predicted to alter the protein coding sequence. The accuracy of the new gene prediction at the *Ave1* locus (two *Ave1* isoforms and one additional gene model) was additionally tested by showing the expression of the *Ave1c* gene. Two sets of primers (*Ave1* and *Ave1c* fw and rev) amplified bands of the expected sizes, confirming the expression of both genes across various *V. dahliae* growth conditions (Fig. 5d). We also attempted to amplify a specific product from both gDNA and cDNA to confirm the orientation and rule out a transcriptional fusion (Fig. 4d, primers *Ave1* fw + *Ave1c* fw). Consistent with the annotation, the amplification using a gDNA template was successful, while the cDNA template failed to amplify a product. Collectively, these results confirm that LoReAn predicts the most accurate gene models at the *Ave1* locus, including a splice-variant of *Ave1.*

### LoReAn produces the most accurate annotation of a second fungal genome using PacBio Iso-seq reads

The basidiomycete *Plicaturopsis crispa,* mostly known for its wood-degrading abilities, has a relatively complex transcriptome with high levels of exons per gene; 5.6 exons per gene compared to *V. dahliae*’s 2.5 exons per gene [45]. Using the settings identified for the *V. dahliae* genome annotation, nine annotations of the *P. crispa* genome were generated using publicly available short-read Illumina RNA-seq and single-molecule PacBio Iso-seq data [46]. The LoReAn annotations predicted the greatest number of genes, transcripts and exons, while BAP and BAP+ had the greatest number of genes, transcripts and exons exactly matching the reference (Table 2). Likewise, the BAP and BAP+ gene, transcript and exon prediction had the highest accuracy when compared to the reference annotation (Fig. 6a). However, the validity of these results is dependent on the quality of the reference annotation. To better understand the output from the annotations in the absence of a potentially confounding reference, the empirical intron analysis was used. Using this analysis of exact match introns, all four LoReAn-based predictions had the highest accuracy, and were even better than the current public reference (Fig. 6b). These results indicate the LoReAn pipeline produces an improved annotation to the current reference based on the mapped RNA-seq data, and that LoReAn using strand information from the sequencing data provides the most accurate annotation of the *P. crispa* genome.

**Table 2.** Predicted features for *P. crispa* annotation analysis.

| **Pipelines<sup>c</sup>** | **Genes (13,626)<sup>a</sup>** |  | **Transcripts (13,636)<sup>a</sup>** |  | **Exons (76,761)<sup>a</sup>** |  |
| --- | --- | --- | --- | --- | --- | --- |
|  | Total Predicted | Exact Match<sup>b</sup> | Total Predicted | Exact Match<sup>b</sup> | Total Predicted | Exact Match<sup>b</sup> |
| Genemark-ES | 11,396 | 4,426 | 11,396 | 4,426 | 73,930 | 56,206 |
| Augustus | 11,640 | 4,976 | 11,640 | 4,976 | 72,936 | 57,648 |
| BAP | 11,831 | 5,489 | 11,831 | 5,489 | 74,598 | 59,693 |
| BAP+ | 11,583 | 5,485 | 11,583 | 5,485 | 72,683 | 59,045 |
| MAKER2 | 8,602 | 2,477 | 11,196 | 2,477 | 62,312 | 44,201 |
| LoRean_M | 14,690 | 5,114 | 15,760 | 5,114 | 75,158 | 57,921 |
| LoRean | 14,698 | 5,112 | 15,765 | 5,112 | 75,228 | 57,935 |
| LoRean_s_M | 12,821 | 5,118 | 13,931 | 5,118 | 74,514 | 57,773 |
| LoRean_s | 12,828 | 5,132 | 13,943 | 5,132 | 74,512 | 57,749 |

<sup>a</sup>The number of reference genes, transcripts and exons are shown in parentheses.
<sup>b</sup>The Exact match column shows the number of predicted features that have the exact genomic location as the reference feature.
<sup>c</sup>Pipelines: BAP - Broad Annotation Pipeline; BAP+ - modified BAP; LoReAn\_M - LoReAn using masked input genome; LoReAn - LoReAn; LoReAn\_s\_M - LoReAn stranded mode using masked input genome; LoReAn\_s - LoReAn stranded mode.

**Fig. 6.**
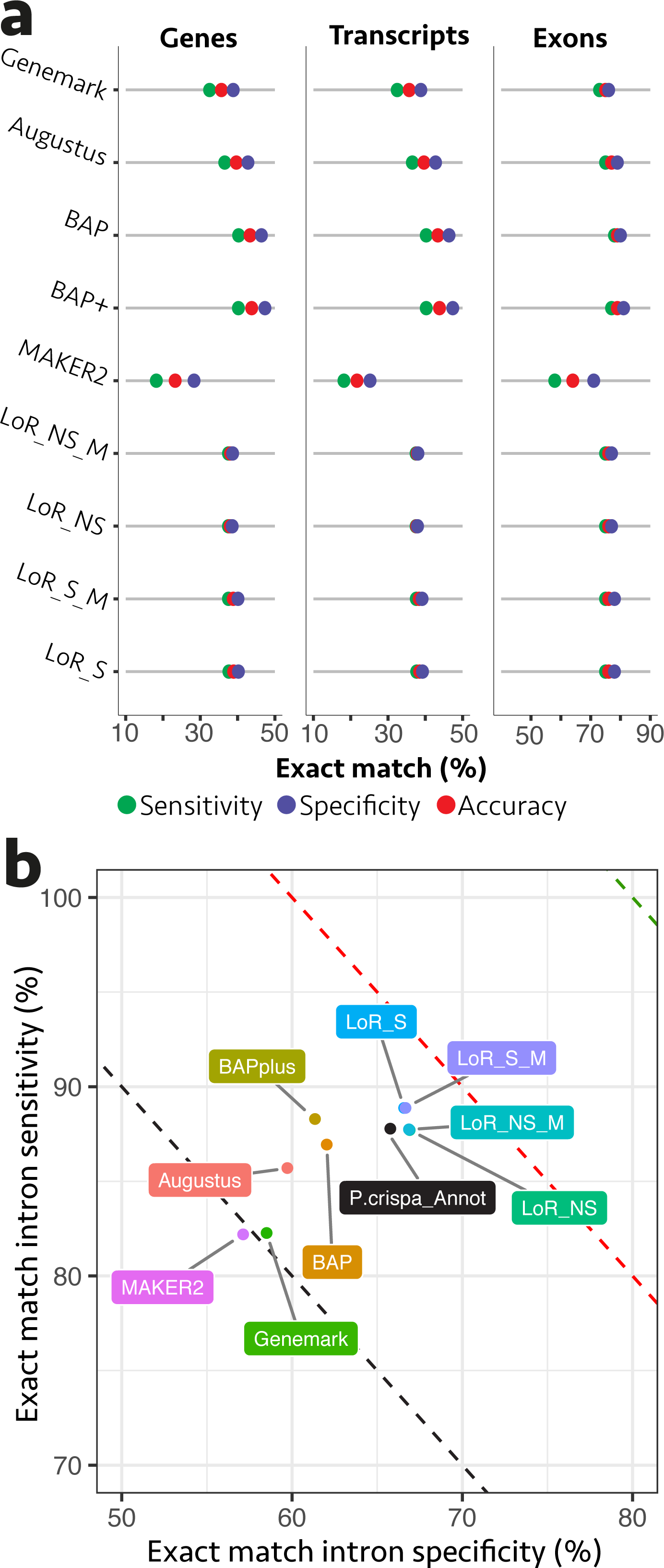
LoReAn gene predictions improve the current *P. crispa* reference annotation. **a** Annotation quality metrics are shown for exact match genes, transcripts and exons labeled at the top of the respective plots. Each horizontal bar represents an annotation output, and each colored dot represents the sensitivity (green), specificity (purple) and accuracy (red). Each output is labeled on the right. LoR_NS_M - LoReAn non-stranded using masked input genome; LoR_NS — LoReAn non-stranded; LoR_S_M — LoReAn stranded using masked input genome; LoR_S - LoReAn stranded. **b** The quality of the annotation pipelines shown was assessed independent of a reference, using the exact match intron location between the gene predictions and those inferred from the short- and long-read mapping data. Sensitivity (y-axis) and specificity (x-axis) were mapped and the average represents their accuracy. The dashed black, red and green lines represent 70%, 80%, and 90% accuracy respectively. Abbreviations are the same as previously detailed. The result from the *P. crispa* reference annotation analysis is shown in black, labeled P. crispa_Annot.

### LoReAn produces high quality annotations for larger plant genomes using PacBio Iso-seq data

To further test LoReAn, the 135 megabase (Mb) *Arabidopsis thaliana* and 375 Mb *Oryza sativa* (rice) genomes were re-annotated using Pacbio Iso-seq data. These genomes are larger and contain a higher percentage of repetitive elements than the two fungal genomes tested. The Arabidopsis annotations generated here were compared to the reference annotation, TAIR10, which is highly curated and represents one of the most complete plant genome annotations [47, 48]. The LoReAn outputs using a non-masked genome had the highest number of genes and transcripts exactly matching the reference, while BAP+ had the highest number of exact match exons (Table 3). The four LoReAn predictions had the highest exact match accuracy compared to the reference for genes, transcripts, and exons (Fig. 7a) We additionally tested the quality of the annotations using exact intron matches to the mapped reads as described earlier. This analysis also shows that the LoReAn outputs were the most accurate and most closely match the TAIR10 reference annotation (Fig. 7b).

**Table 3.** Predicted features for *Arabidopsis thaliana* annotation analysis

| Pipelines<sup>c</sup> | Genes (27,416)<sup>a</sup> |  | Transcripts (35,386)<sup>a</sup> |  | Exons (147,492)<sup>a</sup> |  |
| --- | --- | --- | --- | --- | --- | --- |
|  | Total Predicted | Exact Match<sup>b</sup> | Total Predicted | Exact Match<sup>b</sup> | Total Predicted | Exact Match<sup>b</sup> |
| Genemark-ES | 31,358 | 13,758 | 31,358 | 14,975 | 173,189 | 118,265 |
| Augustus | 27,954 | 15,288 | 27,954 | 16,797 | 161,197 | 121,196 |
| BAP | 29,640 | 15,341 | 29,640 | 16,825 | 163,015 | 121,083 |
| BAP+ | 29,152 | 17,246 | 29,152 | 18,993 | 153,341 | 122,826 |
| MAKER2 | 14,881 | 8,424 | 15,138 | 9,554 | 99,905 | 88,676 |
| LoReAn_M | 24,665 | 17,053 | 25,302 | 19,036 | 133,837 | 119,641 |
| LoReAn | 29,313 | 17,416 | 29,946 | 19,412 | 152,154 | 122,214 |
| LoReAn_s_M | 24,504 | 17,072 | 25,144 | 19,040 | 133,693 | 119,559 |
| LoReAn_s | 29,145 | 17,419 | 29,782 | 19,405 | 152,105 | 122,230 |

<sup>a</sup>The number of reference genes, transcripts and exons are shown in parentheses.
<sup>b</sup>The Exact match column shows the number of predicted features that have the exact genomic location as the reference feature.
<sup>c</sup>Pipelines: BAP - Broad Annotation Pipeline; BAP+ - modified BAP; LoReAn\_M - LoReAn using masked input genome; LoReAn - LoReAn; LoReAn\_s\_M - LoReAn stranded mode using masked input genome; LoReAn\_s - LoReAn stranded mode.

**Fig. 7.**
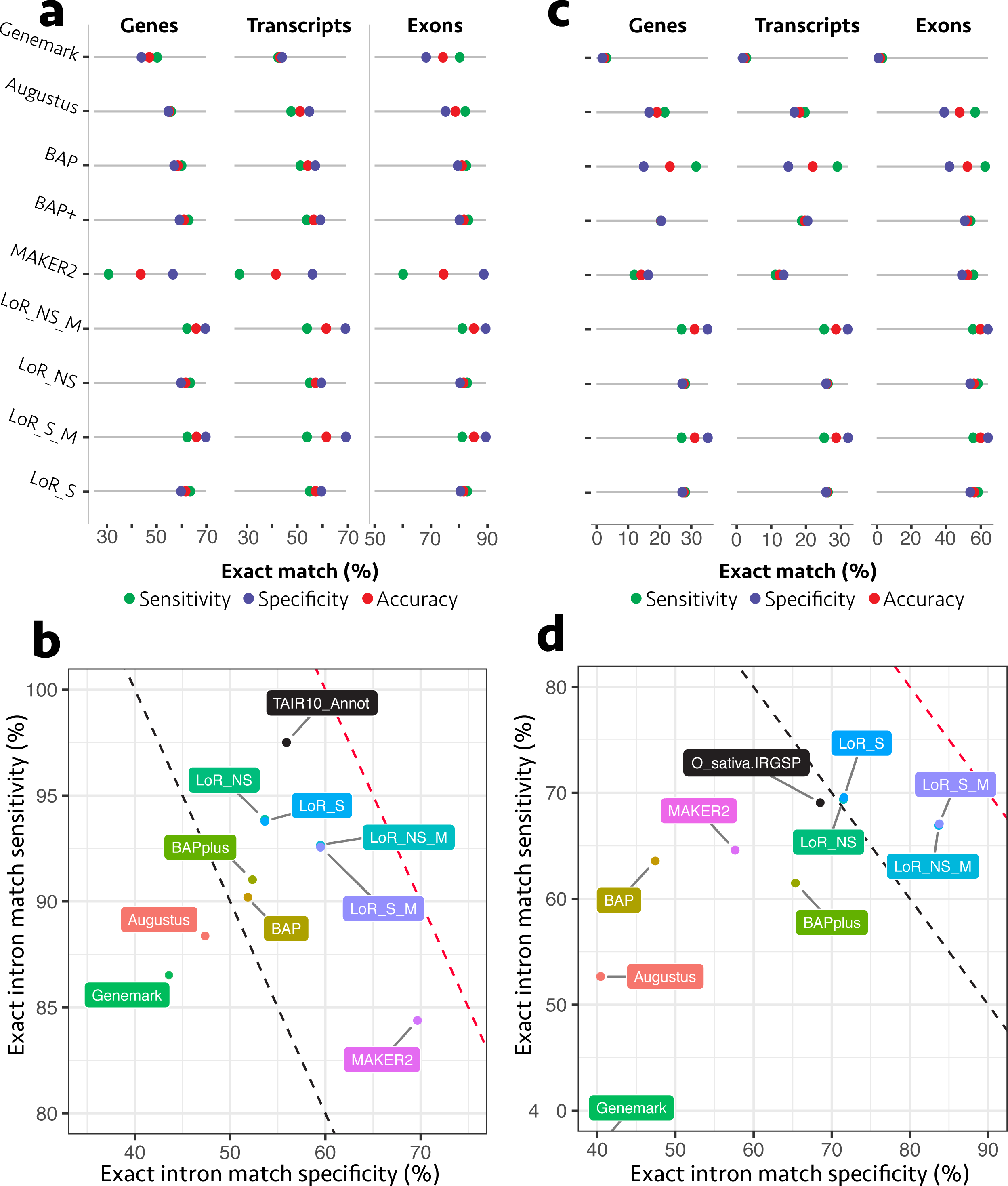
High accuracy LoReAn genome annotations for two plant genomes. **a, c** Annotation quality metrics are shown for exact match genes, transcripts and exons labeled at the top of the respective plots as detailed in figure 6. **a, b** Data for *A. thaliana* **c, d** Data for *O. sativa* **b, d** The quality of the annotation pipelines shown were assessed independent of a reference, using the exact match intron location between the gene predictions and those inferred from the short- and long-read mapping data as detailed in figure 6. **b** The result from the *A. thaliana* reference annotation analysis is shown in black, labeled TAIR10_Annot. **d** The result from the *O. sativa* reference annotation analysis is shown in black, labeled O_sativa.IRGSP.

Comparable results were obtained for the *O. sativa* annotation. The BAP pipeline had the highest number of predicted genes, transcripts and exons exactly matching to the reference annotation, followed by the outputs from the LoReAn predictions (Table 4). However, the four LoReAn predictions had the greatest specificity and accuracy for genes, transcripts and exons compared to the reference annotation (Fig. 7c). The overall level of agreement between the pipelines and the reference is lower for *O. sativa* than for Arabidopsis (compare x-axis, Fig. 7a and 7c), likely reflecting the difference in reference annotation quality. Using the exact intron matches to the mapped reads analysis, the LoReAn gene predictions have the highest accuracy for exact intron matches, 379 even greater than the reference annotation (Fig. 7d). These data suggest LoReAn produced annotations are more accurate than the currently used reference annotation with respect to RNA-seq mapping data.

**Table 4.** Predicted features for *Oryza sativa* annotation analysis.

| **Pipelines<sup>c</sup>** | **Genes (35,679)<sup>a</sup>** |  | **Transcripts (42,132)<sup>a</sup>** |  | **Exons (140,914)<sup>a</sup>** |  |
| --- | --- | --- | --- | --- | --- | --- |
|  | Total Predicted | Exact Match<sup>b</sup> | Total Predicted | Exact Match<sup>b</sup> | Total Predicted | Exact Match<sup>b</sup> |
| Genemark-ES | 62,836 | 1,132 | 62,836 | 1,190 | 512,183 | 4552 |
| Augustus | 46,264 | 7,705 | 46,264 | 8,310 | 205,377 | 79,968 |
| BAP | 75,360 | 11,253 | 75,360 | 12,241 | 209,909 | 88,114 |
| BAP+ | 35,420 | 7,254 | 35,420 | 7,921 | 150,207 | 76,298 |
| MAKER2 | 26,142 | 4,267 | 32,897 | 4,690 | 159,907 | 78,609 |
| LoReAn_M | 27,543 | 9,686 | 32,296 | 10,648 | 122,167 | 78,292 |
| LoReAn | 37,365 | 10,134 | 41,846 | 11,102 | 152,682 | 82,217 |
| LoReAn_s_M | 27,251 | 9,609 | 31,998 | 10,649 | 122,215 | 78,397 |
| LoReAn_s | 37,024 | 10,037 | 41,516 | 11,107 | 152,760 | 82,362 |

<sup>a</sup>The number of reference genes, transcripts and exons are shown in parentheses.
<sup>b</sup>The Exact match column shows the number of predicted features that have the exact genomic location as the reference feature.
<sup>c</sup>Pipelines: BAP - Broad Annotation Pipeline; BAP+ - modified BAP; LoReAn\_M - LoReAn using masked input genome; LoReAn - LoReAn; LoReAn\_s\_M - LoReAn stranded mode using masked input genome; LoReAn\_s - LoReAn stranded mode.

## Discussion

High throughput sequencing continues to have profound impacts on biological systems and the questions researchers are addressing. The technical improvements and associated reduction in cost have resulted in a deluge of high quality model and non-model genomes from across the kingdoms of life. To capture the value of these assembled genomes, equal advances are needed in defining the functional elements of the genome. One such technical advance is the ability to sequence full-length single-molecule cDNAs that directly contain information on transcript structure and alternative forms. This information has previously helped identify alternatively spliced transcripts [26, 49], but singlemolecule long-reads have not been systematically incorporated into annotation pipelines. The newly developed LoReAn pipeline integrates both short-read RNA-seq and long-read single-molecule cDNA sequencing with *ab initio* gene prediction to generate high accuracy gene predictions. In total, three separate analyses using a reference annotation, head-to-head comparison, or comparison to empirical data indicate that LoReAn produces the highest quality annotations of the four genomes tested. These results show that LoReAn has improved performance for predicting gene structures.

Whereas several genome annotation tools use experimental data (i.e. RNA-seq) for gene prediction, none of them fully utilize this information. This is apparent for genes such as *Ave1*, where there is ample RNA-seq evidence supporting the gene model, but most of the tested software fail to predict the gene. This result may be related to the small size of the *Ave1* transcript and the lack of homologs present in fungal databases. The ability to correctly annotate genes with unique features or restricted taxonomic distribution is relevant to many biological questions and will aid comparative genomic studies. We designed LoReAn to provide more weight and incorporate more information from both short- and long-read RNA-seq data as we believe with increasing sequencing depth, length and accuracy this significant source of empirical evidence will greatly improve gene prediction.

The technical and biological characteristics of a genome impacts the annotation options that will influence annotation quality. Genome masking significantly affected the gene prediction output of the *V. dahliae* annotation. From a technical aspect, genome masking prior to annotation likely has the greatest impact when annotating highly contiguously assembled genomes. Fragmented genome assemblies often lack repetitive regions and are *de facto* masked. Masking the telomere-to-telomere complete *V. dahliae* strain JR2 genome resulted in gene predictions which were fragmented because of coding regions overlapping masked regions. Our results indicate that genome masking of short repetitive DNA decreases the quality of the genome annotation, and that using a partial- or non-masked genome may improve annotation results when using long-read data. From a biological perspective, our results show that strand information had a significant impact on annotation quality for the two smaller, more compact fungal genomes. Compact fungal genomes have genes with overlapping UTRs which make gene prediction difficult. Using strand information, LoReAn can assign transcripts to the correct coding strand and avoid the prediction of fused genes. Additionally, strand information is used to assign single exon genes to the correct strand. These results need to be confirmed on a greater number of genomes with diverse characteristics before being fully generalizable. Collectively, our results suggest that both technical and biological information, such as assembly completeness, coding sequence overlap, and intron number per coding sequence impact genome annotation quality and should be considered early during project design.

Our results show that LoReAn can successfully use single-molecule cDNA sequencing data from different platforms to produce high-quality genome annotations, similar to or better than the current community references for four diverse genomes. This shows that the LoReAn pipeline can effectively use single-molecule cDNA sequencing data across the current sequencing platforms and performs well for annotating a small fungal genome of 35 Mb to the rice genome of ~375 Mb. We speculate that the use of annotation software such as LoReAn that incorporates single-molecule cDNA sequencing into the annotation process will significantly improve genome annotation and aid in answering biological questions across all domains of life.

## Conclusions

We present the automated genome annotation software <u>Lo</u>ng <u>Re</u>ad <u>An</u>notation (LoReAn) that builds on previous annotation software to incorporate both short- and long-read sequencing data. This pipeline is shown to perform well using both Oxford Nanopore and Pacific Biosciences produced long-reads and for annotation projects ranging from compact fungal genomes to larger more complex plant genomes. As more labs utilize single-molecule cDNA sequencing to address their specific biological questions, LoReAn will provide an efficient and effective automated annotation pipeline for diverse projects.

## Methods

### Growth conditions and RNA extraction

*Verticillium dahliae* strain JR2 [2], was maintained on potato dextrose agar (PDA) plates grown at approximately 22°C and stored in the dark. Conidiospores were collected from two-week-old PDA plates using half-strength potato dextrose broth (PDB), and subsequently 1×10^6^ spores were inoculated into glass flasks containing 50 mL of either PDB, half-strength Murashige and Skoog (MS) medium supplemented with 3% sucrose, or xylem sap collected from greenhouse grown tomato plants of the cultivar Moneymaker. The cultures were grown for four days in the dark at 22°C and 160 RPM. The cultures were strained through miracloth (22 μm) (EMD Millipore, Darmstadt, Germany), pressed to remove liquid, and flash frozen in liquid nitrogen. Next, the cultures were to ground to powder with a mortar and pestle using liquid nitrogen to ensure samples remained frozen.

RNA extraction was carried out using TRIzol (Thermo Fisher Science, Waltham, MA, USA) following manufacturer guidelines. Following RNA resuspension, contaminating DNA was removed using the TURBO DNA-free kit (Ambion, Thermo Fisher Science, Waltham, MA, USA) and the RNA was checked for integrity by separating 2 μL of each sample on a 2% agarose gel. RNA samples were quantified using a Nanodrop (Thermo Fisher Science, Waltham, MA, USA) and stored at −80°C.

### Library preparation and sequencing — Illumina

Each RNA sample from *V. dahliae* strain JR2 grown in PDB, half-strength MS, and xylem sap was used to construct an Illumina sequencing library for RNA-sequencing by the Beijing Genomics Institute (BGI) following manufacturer guidelines (Illumina Inc., San Diego, CA, USA). Briefly, messenger RNA (mRNA) was enriched using oligo(dT) magnetic beads. The RNA was then fragmented and double stranded cDNA synthesized following manufacturer guidelines (Illumina Inc., San Diego, CA, USA). The fragments were then end-repaired and poly-adenylated to allow for the addition of sequencing adapters, followed by fragment enrichment using polymerase chain reaction (PCR) amplification. Library quality was assessed using the Agilent 2100 Bioanalyzer (Agilent Technologies, Santa Clara, CA, USA). Qualified libraries were sequenced on an Illumina HiSeq-2000 (Illumina Inc., San Diego, CA, USA) at the Beijing Genomics Institute.

### cDNA synthesis and normalization, library preparation and sequencing — Oxford Nanopore Technologies

For the synthesis of single-stranded cDNA, 1 μg of each RNA sample was reverse-transcribed using the Mint-2 cDNA synthesis kit as described by the manufacturer (Evrogen, Moscow, Russia), using the primers PlugOligo-1 (5’ end) and CDS-1 (3’ end). For each sample, 1 μl of cDNA was amplified with PCR for 15 cycles (95°C for 15 seconds, 66°C for 20 seconds and 72°C for 3 minutes) to generate double-stranded cDNA, and purified with 1.8× volume Agencourt AMPure XP magnetic beads (Beckman Coulter Inc., Indianapolis, IN, USA).

Three cDNA samples were normalized with the Trimmer-2 cDNA normalization kit following the manufacturer’s guidelines (Evrogen, Moscow, Russia). The cDNA was precipitated, denatured and hybridized for 5 hours. Next, the double stranded cDNA fraction was cleaved and the remaining single stranded cDNA amplified with PCR for 18 cycles (95°C for 15 seconds, 66°C for 20 seconds and 72°C for 3 minutes).

Library preparation for the three samples was performed using the Nanopore Sequencing Kit (v. SQK-MAP006) following the manufacturer’s guidelines (Oxford Nanopore Technologies [ONT], Oxford, UK). The cDNA was end-repaired and dA-tailed using the NEBNext End Repair and NEBNext dA-Tailing Modules following the manufacturer’s instructions (New England BioLabs [NEB], Ipswich, MA, USA). The reactions were cleaned using an equal volume of Agencourt AMPure XP magnetic beads (Beckman Coulter Inc., Indianapolis, IN, USA), followed by ONT adapter ligation using Blunt/TA ligation Master Mix (NEB, Ipswich, MA, USA). The adapter-ligated fragments were purified using Dynabeads MyOne Streptavidin C1 (Thermo Fisher Science, Waltham, MA, USA).

Sequencing was performed on three different MinION flow cells (v. FLO-MAP103, ONT, Oxford, UK). After priming the flow cells with sequencing buffer, 6 μl of the library preparation was added. Additional library preparation (6 μl) was added to the flow cells at 3, 17 and 24 hours after the run was started. Base-calling was performed using the Metrichor app (v. 2.39.1, ONT, Oxford, UK) and Poretools (v. 0.5.1) was used to generate FASTQ files from the Metrichor produced FAST5 files [50].

### Software in LoReAn pipeline

LoReAn is implemented in Python3. Usage and parameters to run LoReAn, including default settings are detailed at https://github.com/lfaino/LoReAn/blob/master/OPTIONS.md. Mandatory parameters are protein sequences of related organisms, a reference genome sequence and an identification name for the species form the Augustus database. Other inputs are: short-reads (i.e. Illumina RNA-seq) which may be single or paired-end; and long-reads from either MinION or SMRT sequencing platforms. LoReAn outputs a GFF3 file with genome annotations.

The most convenient way to install and run LoReAn is by using the Docker (https://www.docker.com/) image. Information about the software and how to use it can be found at https://github.com/lfaino/LoReAn repository. LoReAn uses the following programs and versions: for read mapping, STAR (version 2.5.3a) [51] and GMAP (v. 2017-06-20) [33]; to assemble and reconstruct transcripts from short reads, Trinity (v. 2.2.0) [32] ran on “genome-guided mode”, followed by PASA (v. 2.1.0) [20]; to map protein sequences, AAT is utilized (v. 03-05-2011) [52]; for gene prediction GeneMark-ES (v4.34) [53] and Augustus (v3.3) [31] are used as *ab initio* software; BRAKER1 (v. 2) [10] is used in substitution of Augustus to generate *ab initio* gene prediction for organism not present in the Augustus catalogue when RNA-seq is supplied; GMAP (v. 2017-06-20) [33] is used for long reads mapping and for assembled ESTs after Trinity assembly; Evidence Modeler (EVM, v. 1.1.1) [20] is used to combine the output from the previous tools to generate a combined annotation model. To extract the genomic sequence, merge and cluster the long-reads, Bedtools suite (v. 2.21.0) [54] is used. iAssembler (v. 1.32) [55] calls a consensus on the clusters (i.e. the process of transcript reconstruction). GenomeTools (v. 1.5.9) software is used at several stages in the LoReAn pipeline [56]. Additional informations about the tools used can be found at https://github.com/lfaino/LoReAn/blob/master/README.md.

### Genome Masking

To study the effect of genome masking on automated genome annotation with LoReAn, we ran the pipeline on stranded mode using three reference genomes with different levels of repetition masking: a fully masked genome with all repetitive sequences masked, a partially masked genome where only repetitions larger than 400 base pairs (bps) were masked and a full genome with no repetition masking. Repeats were masked using RepeatMasker software as previously described [57].

### LoReAn Stranded Mode

To use the software in strand mode efficiently, sequences from the same transcript need to have the same strand. However, sequencing is random and, depending from which fragment and sequencing starts, we can have fragments from the same transcript sequenced in forward or reverse orientation compared to the transcription direction. Unlike DNA sequencing, in cDNA long-read sequencing, the direction of the sequencing can be inferred by localizing only one between the 3’ adapter or the 5’ adapter used during the cDNA production or localizing both. Using the Smith-Waterman alignment, we can identify the location of the adapter/s in the sequenced fragments and adjust the sequencing orientation based on the adapter alignment onto the fragments. For the MinION data we generated, we used the 5’ PlugOligo-1 AAGCAGTGGTATCAACGCAGAGTACGCGGG and 3’-CDS AAGCAGTGGTATCAACGCAGAGTACTGGAG primer sequences associated with the cDNA synthesis and normalization process to identify the coding strand for each long read. For PacBio *Arabidopsis thaliana* experiment, we used the primers AAGCAGTGGTATCAACGCAGAGTACGCGGG and the primer AAGCAGTGGTATCAACGCAGAGTACTTTTT for the correction of the transcript orientation. *Oryza sativa* and *Plicaturopsis crispa* PacBio transcripts were oriented by using the sequence AAAAAAAAAAAAAAAAAAAAAAAAAAAAGTACTCTGCGTTGATACCACTGCTT.

### Annotation quality definitions

We utilized the common metrics sensitivity, specificity and accuracy to compare the annotation features. These metrics have been previously discussed in the context of annotations [7]. Briefly, Sensitivity is a measure of how well an annotation identifies the known features of a reference, also called a true positive rate. For our comparisons, sensitivity can be represented as [(Annotation matching reference / total Reference) * 100] for a specific feature of interest and represents the percentage of known reference features captured. Specificity is a measure of how many of the annotated features are in the reference, also called positive predictive value. For our comparisons, specificity can be represented as the [(Annotation matching reference / total Annotation) * 100] for a specific feature of interest and represents the percentage of all the annotation features that match the reference. These comparisons can be for any annotation feature such as genes, transcripts, or individual exons for exact matches or for a specified overlap to a reference. Accuracy takes both sensitivity and specificity into account and can be represented as [(Sensitivity + Specificity) / 2].

### Head to head comparisons between annotations

To determine the unique protein coding genes annotated between LoReAn-sF, BAP-F, MAKER2 and CodingQuarry we compared the annotations using orthoMCL [40]. OrthoMCL was downloaded from https://github.com/apetkau/orthomcl-pipeline and run using default settings.

### Intron analysis

Introns were extracted from mapped reads using the same methodology from BRAKER1 [10]. Introns supported from at least two reads were extracted and used in the intron set. Genome tool software [56] was used to annotate introns in the gff3 file. Custom scripts were used to identify exact match intron coordinates from the annotation files were overlapped to the intron coordinates from the RNA-seq data. Sensitivity, specificity and accuracy were calculated as described before.

### *Ave1* isoform analysis

*Ave1* isoforms were confirmed using cDNA-PCR of infected plant material with *V. dahliae* strain JR2. Specific primer for the *Ave1* gene (F-TTTAACACTTCACTCTGCTCTCG; R-CCTTGTGTGCTGCTTTGGTA) and for *Ave1c* gene (F-CGCCGGCAATACTATCTCAA; R-ATCCTGTGGGCAACAATAGC) were used to identify the two *Ave1* isoforms. The two forward primers were used to confirm a genomic amplification product, but to disprove a cDNA fusion.

## DECLARATIONS

### Acknowledgements

We thank Jordi Coolen for his assistance in writing the LoReAn software.

### Availability of data and materials

The LoReAn source code is available at: https://github.com/lfaino/LoReAn/ and provided under an MIT license, available at: https://github.com/lfaino/LoReAn/blob/master/LICENSE. Documentation is available at https://github.com/lfaino/LoReAn. The software can run on all platforms when deployed via Docker (https://www.docker.com/).

The *V. dahliae* strain JR2 reference annotation version 5 was used in the analysis. The version 5 was generated by comparing the concordance of all gene models of version 4 with the long reads information. Subsequently, the improved version 5 was deposited at ENSEMBL fungi database and can be downloaded at http://fungi.ensembl.org/Verticilliumdahliaejr2/Info/Index.

The *P. crispa* reference genome and annotation were downloaded from JGI (http://genome.jgi.doe.gov/pages/dynamicOrganismDownload.jsf?organism=Plicr1). The Arabidopsis genome sequence and reference annotation were downloaded from the TAIR database (ftp://ftp.arabidopsis.org/home/tair/Sequences/wholechromosomes/; https://www.arabidopsis.org/downloadfiles/Genes/TAIR10genomerelease/TAIR10gff3/TAIR10GFF3genes.gff). The rice genome sequence and annotation were retrieved from the ENSEMBL plant database (http://plants.ensembl.org/Oryzasativa/Info/Index). The sequencing data are accessible at the NCBI SRA database. The short-read *A. thaliana* data set is deposited under SRA accession number SRR5446746 and the PacBio dataset under SRA accession number SRR5445910. The *V. dahliae* Illumina transcriptome is deposited under accession number SRR5440696 while the Nanopore transcriptome data is deposited as SRR5445874. The *P. crispa* PacBio reads were downloaded from the publicly accessible NCBI SRA site, runs SRR5077068 to SRR5077144 and Illumina data from run SRR1577770. The *O. sativa* data were downloaded from the European Nucleotide Archive (ENA) under runs ERR91110 and ERR911111 and the Illumina data from run ERR748773. All genome annotations, scripts and additional files generated and/or analyzed in the paper can be found at https://github.com/lfaino/files-paper-LoReAn.git. A dataset to test the correct installation of the tool can be found at https://github.com/lfaino/LoReAnExample.git. This dataset contains all the data to annotate a single chromosome of *V. dahliae* strain JR2.

### Authors’ contributions

LF and BT conceived the project. DEC performed data collection for the Illumina sequencing and JVI performed the cDNA normalization and sequencing on the Minion with help from DEC. AP performed the Arabidopsis short- and long-read experiments. HR performed experiments to confirm the annotation results for the *Ave1* locus. LF and JVI wrote the LoReAn python script. LF ran the annotations and LF and DEC performed the analysis. DEC wrote the paper with LF and BT. Funding, guidance and oversight of the project were provided by BT.

### Ethics approval and consent to participate

Ethics approval is not applicable for this study.

## Additional files

Additional file 1: This file contains additional text and Figures S1-S5

Additional file 2: This file contains Tables S1-S4

## References

1. Koren S, Phillippy AM. One chromosome, one contig: complete microbial genomes from long-read sequencing and assembly. Curr. Opin. Microbiol. 2015;23:110–20.

2. Faino L, Seidl MF, Datema E, van den Berg GCM, Janssen A, Wittenberg AHJ, et al. Single-Molecule Real-Time Sequencing Combined with Optical Mapping Yields Completely Finished Fungal Genome. mBio. American Society for Microbiology; 2015;6:e00936-15.

3. Chin C-S, Peluso P, Sedlazeck FJ, Nattestad M, Concepcion GT, Clum A, et al. Phased diploid genome assembly with single-molecule real-time sequencing. Nat. Methods. 2016;13:1050–4.

4. Jiao W-B, Schneeberger K. The impact of third generation genomic technologies on plant genome assembly. Current Opinion in Plant Biology. 2017;36:64–70.

5. Davey JW, Chouteau M, Barker SL, Maroja L, Baxter SW, Simpson F, et al. Major Improvements to the Heliconius melpomene Genome Assembly Used to Confirm 10 Chromosome Fusion Events in 6 Million Years of Butterfly Evolution. G3 (Bethesda). 2016;6:695–708.

6. Thomma BPHJ, Seidl MF, Shi-Kunne X, Cook DE, Bolton MD, van Kan JAL, et al. Mind the gap; seven reasons to close fragmented genome assemblies. Fungal Genet. Biol. 2016;90:24–30.

7. Yandell M, Ence D. A beginner’s guide to eukaryotic genome annotation. Nature Reviews Genetics. Nature Publishing Group; 2012;13:329–42.

8. Cantarel BL, Korf I, Robb SMC, Parra G, Ross E, Moore B, et al. MAKER: an easy-to-use annotation pipeline designed for emerging model organism genomes. Genome Res. Cold Spring Harbor Lab; 2008;18:188–96.

9. Goodswen SJ, Kennedy PJ, Ellis JT. Evaluating high-throughput ab initio gene finders to discover proteins encoded in eukaryotic pathogen genomes missed by laboratory techniques. Tramontano A, editor. PLoS ONE. Public Library of Science; 2012;7:e50609.

10. Hoff KJ, Lange S, Lomsadze A, Borodovsky M, Stanke M. BRAKER1: Unsupervised RNA-Seq-Based Genome Annotation with GeneMark-ET and AUGUSTUS. Bioinformatics. Oxford University Press; 2016;32:767–9.

11. Wang Z, Gerstein M, Snyder M. RNA-Seq: a revolutionary tool for transcriptomics. Nature Reviews Genetics. Nature Publishing Group; 2009;10:57–63.

12. Smith CD, Zimin A, Holt C, Abouheif E, Benton R, Cash E, et al. Draft genome of the globally widespread and invasive Argentine ant (Linepithema humile). Proc. Natl. Acad. Sci. U.S.A. National Acad Sciences; 2011;108:5673–8.

13. Amemiya CT, Alföldi J, Lee AP, Fan S, Philippe H, Maccallum I, et al. The African coelacanth genome provides insights into tetrapod evolution. Nature. Nature Research; 2013;496:311–6.

14. Smith JJ, Kuraku S, Holt C, Sauka-Spengler T, Jiang N, Campbell MS, et al. Sequencing of the sea lamprey (Petromyzon marinus) genome provides insights into vertebrate evolution. Nature Genetics. Nature Research; 2013;45:415-21-421e1-2.

15. Ming R, VanBuren R, Wai CM, Tang H, Schatz MC, Bowers JE, et al. The pineapple genome and the evolution of CAM photosynthesis. Nature Genetics. Nature Research; 2015;47:1435–42.

16. Lamichhaney S, Fan G, Widemo F, Gunnarsson U, Thalmann DS, Hoeppner MP, et al. Structural genomic changes underlie alternative reproductive strategies in the ruff (Philomachus pugnax). Nature Genetics. Nature Research; 2016;48:84–8.

17. Muñoz JF, Gauthier GM, Desjardins CA, Gallo JE, Holder J, Sullivan TD, et al. The Dynamic Genome and Transcriptome of the Human Fungal Pathogen Blastomyces and Close Relative Emmonsia. Haridas S, editor. PLoS Genet. Public Library of Science; 2015;11:e1005493.

18. Linde J, Duggan S, Weber M, Horn F, Sieber P, Hellwig D, et al. Defining the transcriptomic landscape of Candida glabrata by RNA-Seq. Nucleic Acids Res. Oxford University Press; 2015;43:1392–406.

19. Ma L, Chen Z, Huang DW, Kutty G, Ishihara M, Wang H, et al. Genome analysis of three Pneumocystis species reveals adaptation mechanisms to life exclusively in mammalian hosts. Nature Communications. Nature Publishing Group; 2016;7:10740.

20. Haas BJ, Salzberg SL, Zhu W, Pertea M, Allen JE, Orvis J, et al. Automated eukaryotic gene structure annotation using EVidenceModeler and the Program to Assemble Spliced Alignments. Genome Biol. BioMed Central; 2008;9:R7.

21. Haas BJ, Zeng Q, Pearson MD, Cuomo CA, Wortman JR. Approaches to Fungal Genome Annotation. Mycology. Taylor & Francis; 2011;2:118–41.

22. Testa AC, Hane JK, Ellwood SR, Oliver RP. CodingQuarry: highly accurate hidden Markov model gene prediction in fungal genomes using RNA-seq transcripts. BMC Genomics. BioMed Central; 2015;16:170.

23. Phillippy AM. New advances in sequence assembly. Genome Res. Cold Spring Harbor Lab; 2017;27:xi–xiii.

24. Minoche AE, Dohm JC, Schneider J, Holtgräwe D, Viehöver P, Montfort M, et al. Exploiting single-molecule transcript sequencing for eukaryotic gene prediction. Genome Biol. BioMed Central; 2015;16:184.

25. Wang B, Tseng E, Regulski M, Clark TA, Hon T, Jiao Y, et al. Unveiling the complexity of the maize transcriptome by single-molecule long-read sequencing. Nature Communications. Nature Publishing Group; 2016;7:11708.

26. Abdel-Ghany SE, Hamilton M, Jacobi JL, Ngam P, Devitt N, Schilkey F, et al. A survey of the sorghum transcriptome using single-molecule long reads. Nature Communications. Nature Publishing Group; 2016;7:11706.

27. Faino L, Thomma BPHJ. Get your high-quality low-cost genome sequence. Trends in Plant Science. 2014;19:288–91.

28. Laver T, Harrison J, O’Neill PA, Moore K, Farbos A, Paszkiewicz K, et al. Assessing the performance of the Oxford Nanopore Technologies MinION. Biomol Detect Quantif. 2015;3:1–8.

29. Loman NJ, Quick J, Simpson JT. A complete bacterial genome assembled de novo using only nanopore sequencing data. Nat. Methods. 2015;12:733–5.

30. Laehnemann D, Borkhardt A, McHardy AC. Denoising DNA deep sequencing data-high-throughput sequencing errors and their correction. Brief. Bioinformatics. Oxford University Press; 2016;17:154–79.

31. Stanke M, Diekhans M, Baertsch R, Haussler D. Using native and syntenically mapped cDNA alignments to improve de novo gene finding. Bioinformatics. 2008;24:637–44.

32. Grabherr MG, Haas BJ, Yassour M, Levin JZ, Thompson DA, Amit I, et al. Full-length transcriptome assembly from RNA-Seq data without a reference genome. Nature Biotechnology. 2011;29:644–52.

33. Wu TD, Watanabe CK. GMAP: a genomic mapping and alignment program for mRNA and EST sequences. Bioinformatics. Oxford University Press; 2005;21:1859–75.

34. Križanovic K, Echchiki A, Roux J, Šikic M. Evaluation of tools for long read RNA-seq splice-aware alignment. Bioinformatics. 2017.

35. Fradin EF, Thomma BPHJ. Physiology and molecular aspects of Verticillium wilt diseases caused by V. dahliae and V. albo-atrum. Mol. Plant Pathol. Blackwell Publishing Ltd; 2006;7:71–86.

36. Klosterman SJ, Atallah ZK, Vallad GE, Subbarao KV. Diversity, pathogenicity, and management of verticillium species. Annu Rev Phytopathol. Annual Reviews; 2009;47:39–62.

37. Keibler E, Brent MR. Eval: a software package for analysis of genome annotations. BMC Bioinformatics. BioMed Central; 2003;4:50.

38. Chan K-L, Rosli R, Tatarinova TV, Hogan M, Firdaus-Raih M, Low E-TL. Seqping: gene prediction pipeline for plant genomes using self-training gene models and transcriptomic data. BMC Bioinformatics. BioMed Central; 2017;18:1426–7.

39. Chen F, Mackey AJ, Stoeckert CJ, Roos DS. OrthoMCL-DB: querying a comprehensive multi-species collection of ortholog groups. Nucleic Acids Res. 2006;34:D363–8.

40. Li L, Stoeckert CJ, Roos DS. OrthoMCL: identification of ortholog groups for eukaryotic genomes. Genome Res. Cold Spring Harbor Lab; 2003;13:2178–89.

41. Cook DE, Mesarich CH, Thomma BPHJ. Understanding plant immunity as a surveillance system to detect invasion. Annu Rev Phytopathol. Annual Reviews; 2015;53:541–63.

42. Presti Lo L, Lanver D, Schweizer G, Tanaka S, Liang L, Tollot M, et al. Fungal effectors and plant susceptibility. Annu Rev Plant Biol. Annual Reviews; 2015;66:513–45.

43. Sperschneider J, Dodds PN, Gardiner DM, Manners JM, Singh KB, Taylor JM. Advances and challenges in computational prediction of effectors from plant pathogenic fungi. Sheppard DC, editor. PLoS Pathog. Public Library of Science; 2015;11:e1004806.

44. de Jonge R, van Esse HP, Maruthachalam K, Bolton MD, Santhanam P, Saber MK, et al. Tomato immune receptor Ve1 recognizes effector of multiple fungal pathogens uncovered by genome and RNA sequencing. Proc. Natl. Acad. Sci. U.S.A. National Acad Sciences; 2012;109:5110–5.

45. Gordon SP, Tseng E, Salamov A, Zhang J, Meng X, Zhao Z, et al. Widespread Polycistronic Transcripts in Fungi Revealed by Single-Molecule mRNA Sequencing. Zheng D, editor. PLoS ONE. Public Library of Science; 2015;10:e0132628.

46. Kohler A, Kuo A, Nagy LG, Morin E, Barry KW, Buscot F, et al. Convergent losses of decay mechanisms and rapid turnover of symbiosis genes in mycorrhizal mutualists. Nature Genetics. Nature Research; 2015;47:410–5.

47. Lamesch P, Berardini TZ, Li D, Swarbreck D, Wilks C, Sasidharan R, et al. The Arabidopsis Information Resource (TAIR): improved gene annotation and new tools. Nucleic Acids Res. Oxford University Press; 2012;40:D1202–10.

48. Berardini TZ, Reiser L, Li D, Mezheritsky Y, Muller R, Strait E, et al. The Arabidopsis information resource: Making and mining the “gold standard” annotated reference plant genome. Genesis. 2015;53:474–85.

49. Au KF, Sebastiano V, Afshar PT, Durruthy JD, Lee L, Williams BA, et al. Characterization of the human ESC transcriptome by hybrid sequencing. Proc. Natl. Acad. Sci. U.S.A. National Acad Sciences; 2013;110:E4821–30.

50. Loman NJ, Quinlan AR. Poretools: a toolkit for analyzing nanopore sequence data. Bioinformatics. Oxford University Press; 2014;30:3399–401.

51. Dobin A, Davis CA, Schlesinger F, Drenkow J, Zaleski C, Jha S, et al. STAR: ultrafast universal RNA-seq aligner. Bioinformatics. Oxford University Press; 2013;29:15–21.

52. Huang X, Adams MD, Zhou H, Kerlavage AR. A tool for analyzing and annotating genomic sequences. Genomics. 1997;46:37–45.

53. Lomsadze A, Burns PD, Borodovsky M. Integration of mapped RNA-Seq reads into automatic training of eukaryotic gene finding algorithm. Nucleic Acids Res. 2014;42:e119-9.

54. Quinlan AR, Hall IM. BEDTools: a flexible suite of utilities for comparing genomic features. Bioinformatics. Oxford University Press; 2010;26:841–2.

55. Zheng Y, Zhao L, Gao J, Fei Z. iAssembler: a package for de novo assembly of Roche-454/Sanger transcriptome sequences. BMC Bioinformatics. BioMed Central; 2011;12:453.

56. Gremme G, Steinbiss S, Kurtz S. GenomeTools: a comprehensive software library for efficient processing of structured genome annotations. IEEE/ACM Trans Comput Biol Bioinform. 2013;10:645–56.

57. Faino L, Seidl MF, Shi-Kunne X, Pauper M, van den Berg GCM, Wittenberg AHJ, et al. Transposons passively and actively contribute to evolution of the two-speed genome of a fungal pathogen. Genome Res. Cold Spring Harbor Lab; 2016;26:1091–100.

